# Experimental evaluation of environmental DNA detection of a rare fish in turbid water

**DOI:** 10.1101/2022.08.24.502857

**Authors:** Ann E. Holmes, Melinda R. Baerwald, Jeff Rodzen, Brian M. Schreier, Brian Mahardja, Amanda J. Finger

## Abstract

Environmental DNA (eDNA) approaches enable sensitive detection of rare aquatic species. However, water conditions like turbidity can limit sensitivity, resulting in false negative detections. The dynamics of eDNA detection in turbid conditions are poorly understood, but can be better characterized through experimental work. In this study, 1-L field-collected water samples were spiked with tank-sourced eDNA from a rare, endangered estuarine fish at concentrations similar to eDNA samples collected from the natural environment. Samples using non-turbid water (5 NTU), turbid water (50 NTU), and prefiltered turbid water were filtered using four filter types (pore size range 0.45 μm-10 μm). Detection success using a species-specific Taqman qPCR assay was assessed as both eDNA copy number and detection/non-detection. Glass fiber filters (nominal pore size 1.6 μm) yielded the highest number of eDNA copies and detections in non-turbid water and the highest detection rate in turbid water when used without a prefilter. Detection was a more robust metric for evaluating species presence across turbidity conditions compared with eDNA copy number. Prefiltration improved detection rates for the other filters tested (polycarbonate and cartridge filters). Filter material and design appear to interact differently with the prefiltration step, and may be more important considerations than pore size for eDNA capture in turbid water. Interactions between eDNA particles, suspended particulate matter, and filters are important to consider for eDNA methods optimization and interpretation of rare species detections in turbid water.

## Introduction

Environmental DNA (eDNA) can be an efficient tool for surveying species; it can be as or more sensitive than conventional survey methods (Jerde et al. 2011; Shaw et al. 2016; Sigsgaard et al. 2017) and detect species not detected using other methods (e.g., Budd et al. 2021; Renan et al. 2017). Indirect detection using eDNA poses little or no risk of sampling-related mortality or stress to both target and non-target organisms, an advantage when targeting or sampling in the vicinity of endangered or sensitive species. Moreover, eDNA samples can usually be collected with less risk to personnel in potentially hazardous conditions (e.g., high gradient streams and rivers). However, despite the advantages of using eDNA to survey rare species, challenging environmental conditions may adversely affect detection sensitivity, resulting in false negative detections.

The sensitivity of eDNA detection is influenced by interacting suites of biological and environmental conditions (Barnes and Turner 2016). Target organism biomass, individual body size, and eDNA shed rate may influence eDNA detection probability (e.g., Sassoubre et al. 2016). Environmental conditions such as water movement, turbidity, temperature, pH, salinity, solar radiation, or microbial community composition (or the related biotic conditions) also affect species detection (e.g., Collins et al. 2018; Jane et al. 2015; Laramie et al. 2015; Seymour et al. 2018; Shogren et al. 2018; Strickler et al. 2015; Tsuji et al. 2019). Turbidity is a previously recognized challenge for eDNA sampling and a suspected cause of reduced sensitivity and false negative detections (Egeter et al. 2018; Williams et al. 2017).

Turbidity can affect eDNA detection in a variety of ways. Turbidity is a measure of light scatter in water and is associated with reduced water clarity, although they are not necessarily equivalent measurements. Turbidity is caused by a variety of unrelated phenomena including particulates spanning a range of sizes and compositions (e.g., sediment, inorganic material, or organic material such as plankton and plant detritus) and decreased water clarity without particulates (e.g., staining by tannins from plant material). Both particulates and staining can introduce PCR inhibitors (e.g., humic compounds; Matheson et al. 2010) that interfere with molecular detection. The effects of PCR inhibitors can be effectively removed without diluting samples using appropriate extraction methods (Hunter et al. 2015) or with a post-extraction inhibitor removal step (Williams et al. 2017).

Suspended particulate matter remains a major challenge for eDNA detection. Particulates clog filters, leading to decreased filtration volumes and long filtration times. Particulate matter can decrease sensitivity or eliminate eDNA detections altogether even when the target species is present (Day et al. 2019). Species detection sensitivity has been positively correlated with volume of water sampled (Hunter et al. 2019; Schabacker et al. 2020; Sepulveda et al. 2019). Sample volume has been shown to influence eDNA detection sensitivity more than the number of samples or quantitative PCR (qPCR) replicates (Schultz and Lance 2015). However, long filtration times may not be worth the wait; measurements of membrane pressure suggest diminishing returns on eDNA detection due to increased pressure during filtration (Thomas et al. 2018). Filters with larger pores (Robson et al. 2016) and prefiltration (Takahara et al. 2012) are recommended to increase water volume and decrease filtering time in turbid systems. While some results suggest a positive effect of prefiltration on eDNA capture (Takahara et al. 2012), presumably due to increased volume filtered, others are inconclusive (Majaneva et al. 2018). A better understanding of the impact of turbidity and methodological adjustments on eDNA detection is necessary, particularly when a rare species of interest is positively associated with turbidity (e.g., Feyrer et al. 2007; Nobriga 2002; Sommer et al. 2011).

In this study, we examine the effects of turbidity and filtration methods for detection of target eDNA in low concentrations. We tested four filter types (pore size range 0.45 μm-10 μm) and the addition of a prefiltration step for turbid water. Filters were chosen to capture fish mitochondrial eDNA particles in the size range where they are most abundant (1-10 μm; Turner et al. 2014; Wilcox et al. 2015). We hypothesized that (1) turbidity would decrease both eDNA copies detected and the detection rate, (2) larger filter pore sizes would partially offset the negative effects of suspended particulate matter on eDNA detection, and (3) prefiltration would improve detection in turbid water.

## Materials & Methods

### Study species and habitat

Delta smelt (*Hypomesus transpacificus*) are small (5-7 cm), critically endangered fish endemic to the San Francisco Estuary (SFE), California, USA. Delta smelt are considered the sentinel species of the SFE ecosystem (Moyle et al. 2018), significant in indigenous Miwko? (Miwok) traditional cultural practice and law (Hankins 2018), and at risk of extinction in the near future (Moyle et al. 2018). Delta smelt typically have an annual life cycle and are unusually sensitive to changes in estuarine conditions (Moyle et al. 1992). The species is protected under both the Federal Endangered Species Act (ESA) and California Endangered Species Act (CESA) due to a 90% decline in population abundance over the two decades prior to listing in 1993 (USFWS 1993). Around 2000, abundance of delta smelt and other pelagic fishes in the SFE again declined dramatically, most likely due to environmental factors including changes in water quality, habitat degradation, and effects of introduced species (Sommer et al. 2007; Moyle et al. 2016).

The presence of delta smelt is positively associated with turbid water (Feyrer et al. 2007; Nobriga et al. 2008; Sommer et al. 2011) perhaps due to decreased predation risk (Ferrari et al 2014; Bennett and Burau 2015) and increased larval feeding rates (Baskerville-Bridges et al. 2004; Tigan et al. 2020). The physiological performance of delta smelt is negatively affected by turbidity levels below 25 NTU and above 80 NTU (Hasenbein et al. 2016). Turbidity in delta smelt habitat is attributed to suspended sediment transported from upstream sources or resuspended in the water column due to wind or turbulence (Schoellhamer 2002).

Delta smelt are difficult to survey due to extremely low abundance, despite exceptional monitoring efforts by state and federal agencies. The U.S. Fish and Wildlife Service (USFWS) began year-round, spatially extensive surveys targeting delta smelt using multiple (conventional) gear types in late 2016 (Enhanced Delta Smelt Monitoring program (EDSM; USFWS 2022). The EDSM provides data on distribution and abundance of delta smelt and other species of concern for conservation and management (Mahardja et al. 2021). Pilot eDNA surveys of delta smelt conducted alongside EDSM trawls and indicated concordance with trawl sampling, but single positive qPCR replicates for each sample provide weak evidence of species presence (Supplementary File S1; Goldberg et al. 2016). Moreover, already low trawl detection rates continued to decrease (USFWS 2022), making further field testing of eDNA methods unfeasible. Experimental testing was undertaken to determine if turbidity and filtration methods were major constraints on eDNA detection of delta smelt.

### In vivo testing

Quantitative PCR (qPCR) detection of delta smelt eDNA used a Taqman probe and primers previously validated using genomic DNA and tested for cross-reactivity with congener Wakasagi smelt (*Hypomesus nipponensis*) and 21 other SFE fish species (Baerwald et al. 2011). In this study, the Limit of Detection (LOD) and Limit of Quantification (LOQ) were determined following guidelines for standardized analysis of eDNA samples (Klymus et al. 2019; Merkes et al. 2019) using serial dilutions of a synthetic oligonucleotide gBlocks Gene Fragment (Integrated DNA Technologies, San Diego, CA) of a portion of the delta smelt cytochrome b gene assayed in 8 replicates with a starting concentration of 0.1 pg/ul (3.5 x 10^6^ copies/reaction) and 1:4 subsequent dilution.

LOD is defined as the lowest concentration in which the target molecule can be detected in 95% of replicates (Bustin et al. 2009). The theoretical minimum LOD is 3 copies of template DNA per PCR reaction, assuming a Poisson distribution of the target molecules in PCR reactions. The effective LOD is applied to multiple qPCR replicates, showing a decrease in LOD with increasing replicates (Klymus et al. 2019). The LOQ assesses precision using the coefficient of variation (CV) of the measured concentrations of DNA standards (Kubista 2014). The LOQ is defined as the lowest concentration at which the CV of qPCR results is less than 35% (Klymus et al. 2019).

### Filtration experiment

Filtration used a peristaltic pump (Geotech Environmental Equipment Inc., Denver, Colorado). Bottles and other materials used for filtering were sterilized for at least 20 min in 20% bleach then rinsed three times with clean water. Tubing was sterilized by pumping 20% bleach through the tube for at least 60 sec then flushing the tube with clean water for at least 60 sec. The samples were set up and filtered in a laboratory space free from delta smelt tanks, tissue, or DNA.

Estuarine water was collected in sterilized 5-gal buckets from two sites in a freshwater region of the upper SFE where delta smelt are not present (Figure 1). Turbidity of the water collected at the sites was measured at ~5 NTU (“non-turbid”) and ~50 NTU (“turbid”) with a Hach 2100Q portable turbidimeter. The non-turbid and turbid designations are relative measures and ecologically relevant to delta smelt; 50 NTU is less turbid than conditions regularly observed in winter in the SFE (>100 NTU). The buckets were covered and transported to UC Davis campus. Water was homogenized by stirring with a sterilized implement before being transferred to sterilized 1 L bottles for the experiment.

**Figure 1.**
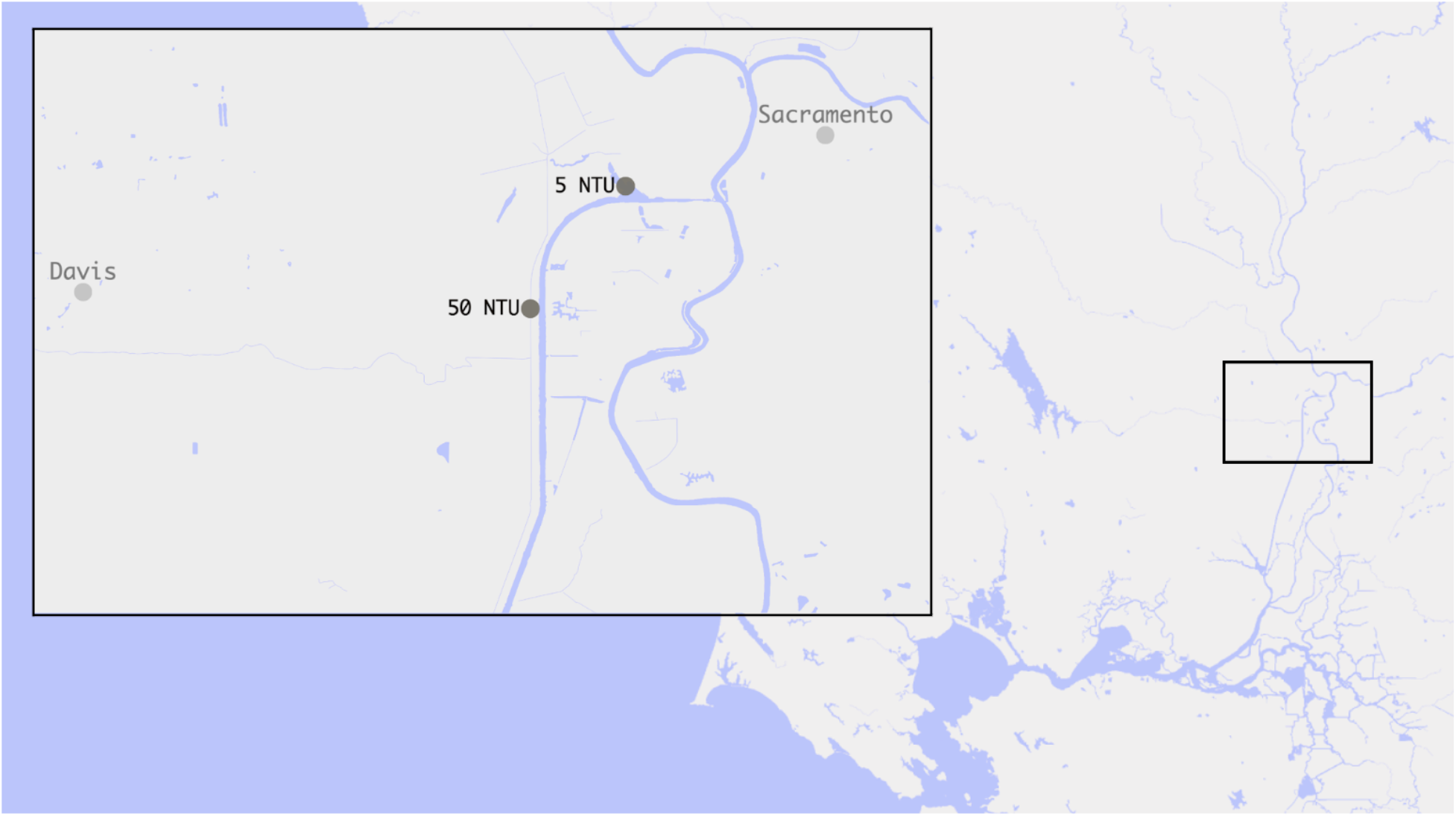
Upper San Francisco Estuary (California, USA) collection sites (inset) for water used in filtration experiments. Non-turbid water (~5 NTU) was collected from the upper Sacramento Deep Water Shipping Channel (38.5653, −121.5539) and turbid water (~50 NTU) was collected from upper Prospect Slough (35.5299, −121.589), adjacent to the shipping channel. (Map made using kepler.gl and mapbox.)

A schematic of the study design is shown in Figure 2. Water from a 340-L tank (recirculating aquaculture system with daily make-up water to maintain a tank volume) containing an estimated 186 adult delta smelt at the UC Davis Center for Aquatic Biology and Aquaculture was collected in a sterile 1-L bottle and stored on wet ice. The bottle was gently inverted several times to homogenize eDNA prior to pipetting 0.5 mL tank water into each 1 L bottle of estuarine water. Pilot experiments determined that 0.5 mL tank water added to 1 L of estuary water produced Ct values similar to field detections. The same process was repeated but 1 mL was added to each bottle in case 0.5 mL tank water was undetectable in some replicates. Adding two small but different volumes of tank water also allowed us to assess whether small differences in eDNA concentration can be distinguished at low concentrations. Bottles were placed in a sterilized cooler with wet ice and filtered within ~8 hours.

**Figure 2.**
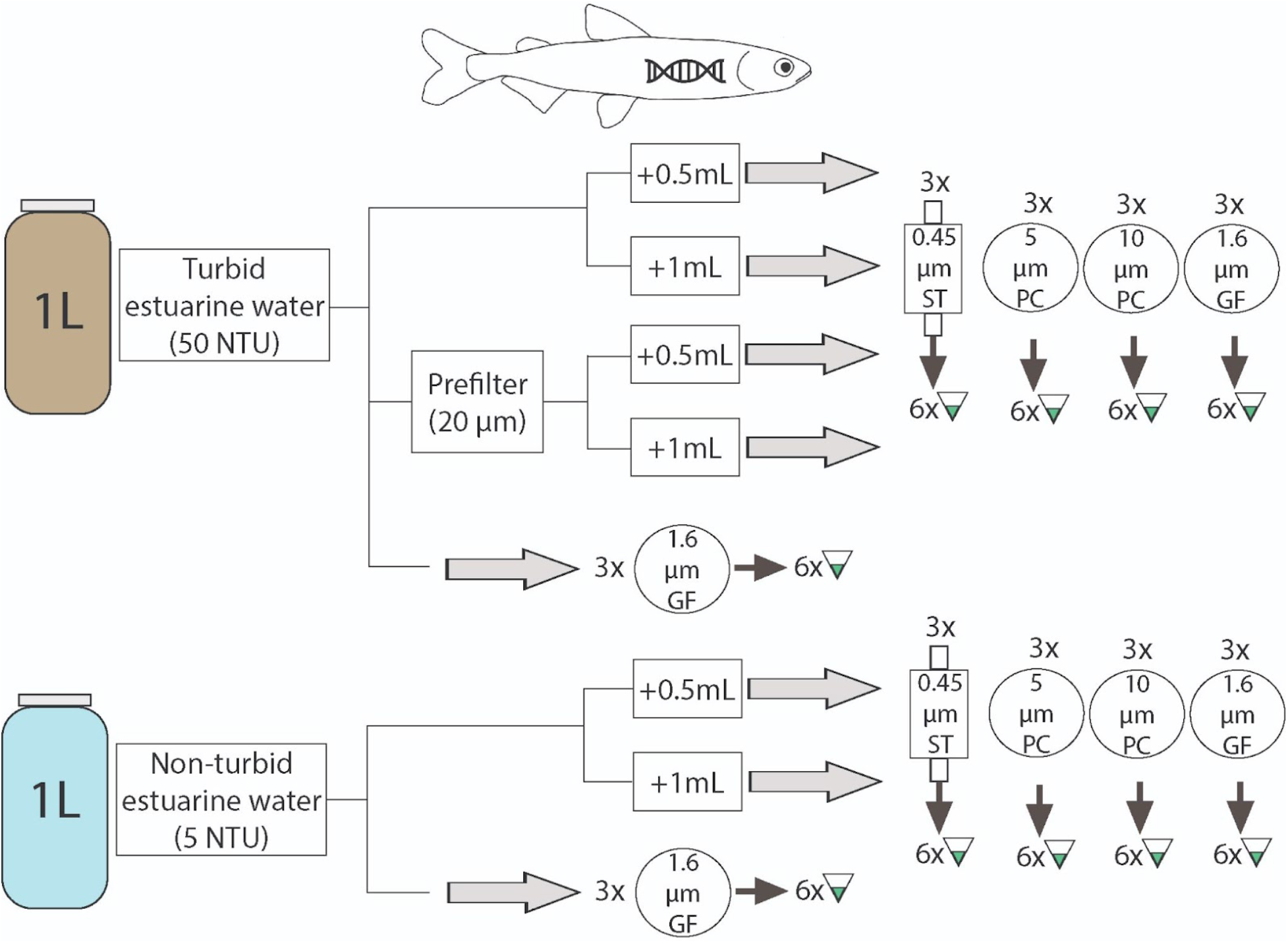
Schematic representation of filtration experiment. Three biological replicates of 6 treatments were each filtered on 4 different filter types and assessed in 6 qPCR replicates using a species-specific assay (Baerwald et al. 2011). GF, glass fiber filter; PC, polycarbonate filter; ST, Sterivex cartridge filter.

Each bottle was gently inverted several times prior to filtering. Three biological replicates were filtered using each of the four filter types in the three treatments: non-turbid water, turbid water, and turbid water with the addition of a prefilter (Table 1). This design resulted in 72 biological replicates (1-L bottles). Glass fiber, polycarbonate filters, and nylon mesh prefilters were loaded into sterile filter holders (Swinnex-47, MilliporeSigma) attached to silicon tubing. Sterivex filter cartridges were attached directly to the tubing. Water was pumped through filters until the 1-L samples was filtered or flow ceased (usually a maximum of ~15 min). Filtration volumes less than 1 L reflect filter clogging in turbid water. After filtration, glass fiber and polycarbonate filters were folded twice and placed in a sterile 2 mL tube and Sterivex filters were capped at each end. The tubes or capped cartridges were placed in individual sterile plastic bags and immediately frozen on dry ice. Frozen samples were transferred to −20°C for storage until extraction. Three negative control samples of estuarine water from each turbidity value without added tank water were processed with the field samples.

**Table 1.** Characteristics of the filters and prefilter used in the filtration experiment.

| Filters | Abbreviation | Filter shape | Filter type | Pore size | Pore type |
| --- | --- | --- | --- | --- | --- |
| Sterivex polyvinylidene fluoride (PVDF; MilliporeSigma) | ST | cartridge | screen | 0.45 µm | absolute |
| Glass fiber (Whatman) | GF | 47mm diameter | depth | 1.6 µm | nominal |
| Polycarbonate track-etched (MilliporeSigma) | PC | 47mm diameter | screen | 5 µm | absolute |
| Polycarbonate track-etched (MilliporeSigma) | PC | 47mm diameter | screen | 10 µm | absolute |
| **Prefilter** |  |  |  |  |  |
| Nylon net (MilliporeSigma) | NN | 47mm diameter | screen | 20 µm | mesh |

Genetics work was conducted in a dedicated eDNA laboratory space following recommended guidelines (Goldberg et al. 2016). DNA was extracted in a dedicated eDNA extraction hood using the DNeasy PowerWater Kit (Qiagen, Hilden, Germany), which has been shown to effectively remove PCR inhibitors (Eichmiller et al. 2015). DNA from whole 47 mm filters was extracted using the DNeasy PowerWater Kit and from Sterivex cartridges were extracted using the DNeasy PowerWater Sterivex Kit. Extraction protocols followed the manufacturer’s instructions including the optional heat lysis step (Supplementary File S2). Elution buffer incubation time was extended to ~20 minutes and DNA was eluted into LoBind tubes (Eppendorf, Hamburg, Germany).

Six technical replicates (PCR reactions) of each sample and the estuary water negative controls were assayed using a species-specific Taqman assay targeting delta smelt (Baerwald et al. 2011). qPCR reactions set-up in a dedicated eDNA PCR hood used TaqMan Environmental Master Mix 2.0 (Applied Biosystems, Waltham, MA, USA). Reagent volumes and cycling conditions are listed in Table 2. qPCR was conducted on a single CFX Touch Real-Time PCR instrument (Bio-Rad Laboratories, Hercules, CA, USA) in a laboratory room separate from eDNA extraction and PCR setup hoods. No-template qPCR controls and gBlock qPCR positive control samples were also assayed using the same protocol.

**Table 2.**
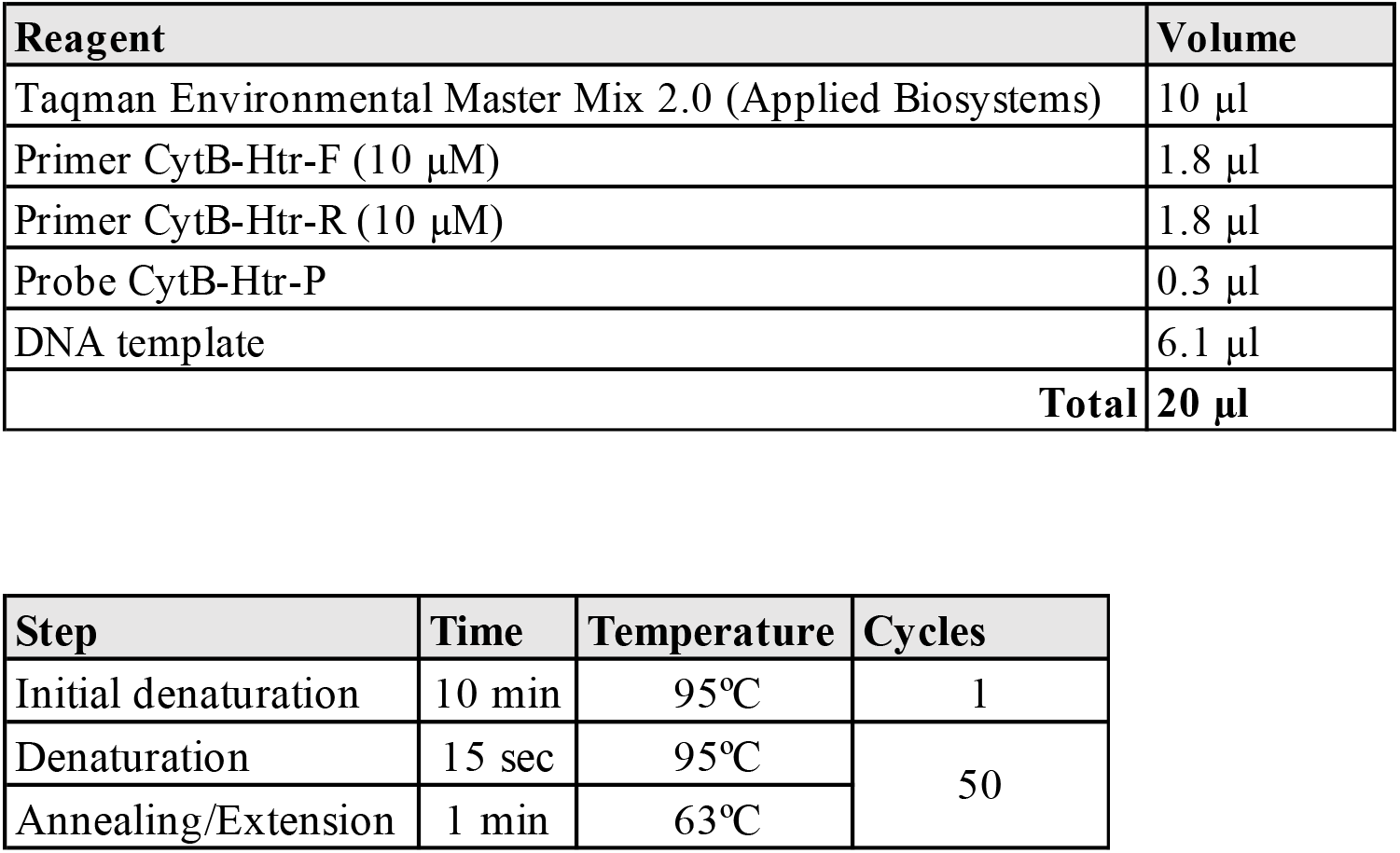
Reagent volumes (a) and qPCR thermocycling protocol (b) for delta smelt eDNA detection.

### Statistical modeling

Results were analyzed using generalized linear mixed-effect models (GLMMs) and a model comparison approach. Analysis was conducted in R (R Core Team, 2017) using packages lme4 (Bates et al. 2015) and bbmle (Bolker and R Development Core Team, 2017). The models used results in non-turbid water, turbid water without a prefilter, and turbid water with a prefilter (n=144 qPCR reactions for each treatment) to identify factors that influence success of delta smelt detection under conditions similar to those observed in the natural environment (turbidity and low eDNA concentrations).

Either eDNA copy number or detection/non-detection can be used to evaluate the factors that influence detection. Copy number allows for a more nuanced interpretation of detection success, but may not be reliable for low eDNA concentrations (i.e., below the LOQ). We modeled both eDNA copies detected (as log(copies+1)) and detection/non-detection in each qPCR replicate as response variables in two models. The full model for both analyses included five covariates as fixed effects: “turbidity” (a categorical variable with two levels corresponding to 5 NTU or 50 NTU); “filter type” (a categorical variable with four levels corresponding to the four filter types); “prefilter” (a categorical variable with two levels), “volume filtered” (a continuous variable of the volume of water filtered rounded to the nearest 50 mL), and “volume of tank water added” (a categorical variable with two levels corresponding to addition of 0.5 or 1 mL of water from the tank of delta smelt). Interactions between both turbidity and prefilter with filter type were also considered. Full models included biological replicate (each 1-L bottle filtered) as a random effect to account for bottle-to-bottle (biological replicate) variation within treatments. Models were compared using Akaike’s Information Criterion corrected for small sample size (AICc).

## Results

### In vivo testing

The one replicate Limit of Detection (LOD) was 2.47 copies per PCR reaction (SE 1.59) and the Limit of Quantification (LOQ) was 67 copies per PCR reaction (Figure 2; Supplementary Files S3, S4). Although less than the theoretical minimum of 3 copies per reaction, the calculated LOD calculation is acceptable because it falls within the calculated error (Bustin et al. 2009). The effective LOD for six qPCR replicates (the number used in this study) was 1.02 copy per PCR reaction (SE 0.15; Supplementary File S4).

### Filtration experiment

In non-turbid water, eDNA detection was 100% for glass fiber, cartridge, and 5-μm pore polycarbonate filters, and close to 100% for 5-μm pore polycarbonate filters (Figure 4; Table 3; Supplemental File S5). eDNA copies were highest for glass fiber filters and cartridge filters, despite the lower volume filtered by cartridge filters due to clogging (Figure 5; Table 3; Supplemental File S5). In turbid water, eDNA detection was nearly 100% for glass fiber filters, but below ~75% for Sterivex filters and at or below 50% for both polycarbonate filters (Figure 4; Table 3; Supplemental File S5). Turbidity reduced the number of eDNA copies detected for all samples, especially the Sterivex and polycarbonate filters (Figure 5; Table 3; Supplemental File S5). The addition of a prefilter increased copy numbers and detection rate for all samples except those collected on glass fiber filters, which were negatively impacted by prefiltration (Figure 4, 5; Table 3; Supplemental File S5). Delta smelt eDNA was not detected in negative controls of estuarine water or qPCR no template controls.

**Figure 3.**
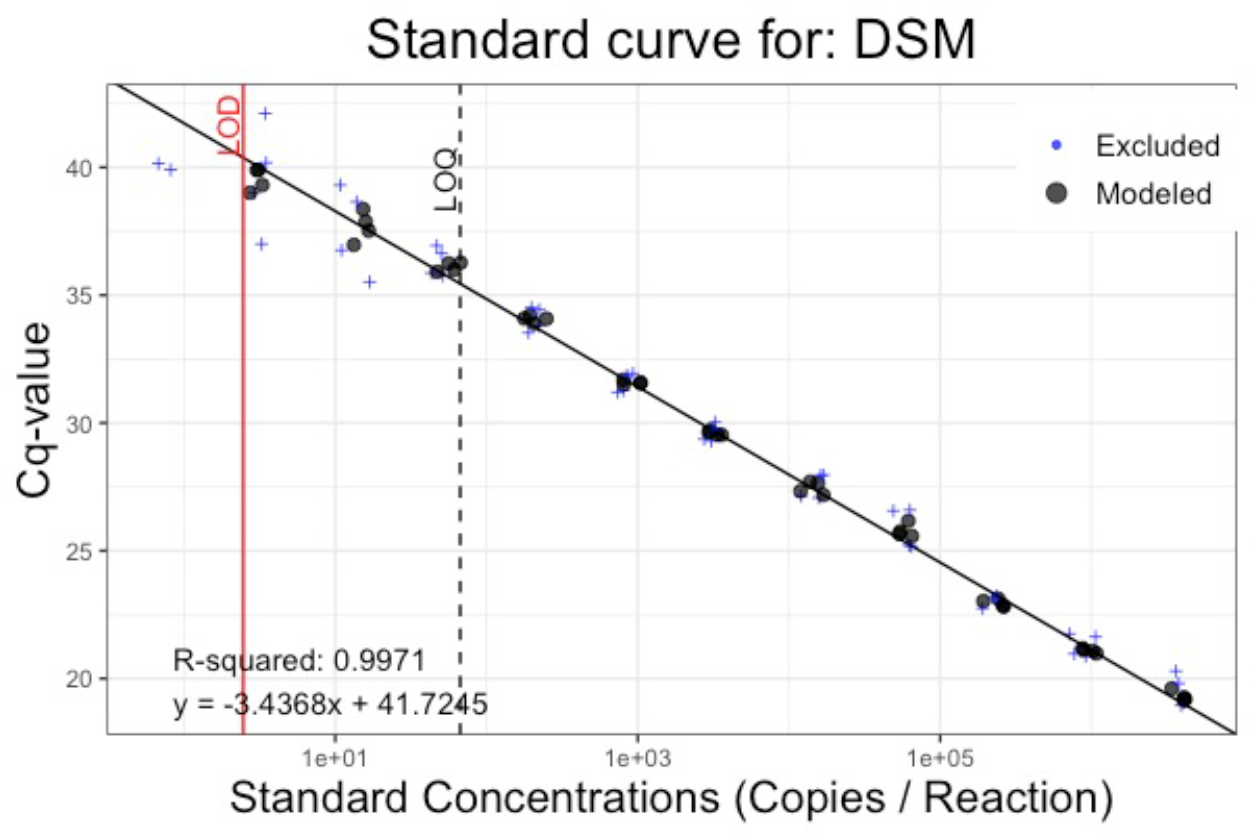
Calibration curve showing Limit of Detection (LOD) for one replicate (2.47 copies) and the Limit of Quantification (LOQ; 67 copies) for the delta smelt Taqman assay (Baerwald et al. 2011). Calculations follow standard methods for validating eDNA assays (Klymus et al. 2019; Merkes et al. 2019). Only points in the middle 2 quartiles of standards with at least 50% detection (black circles) are included in the calculations.

**Figure 4.**
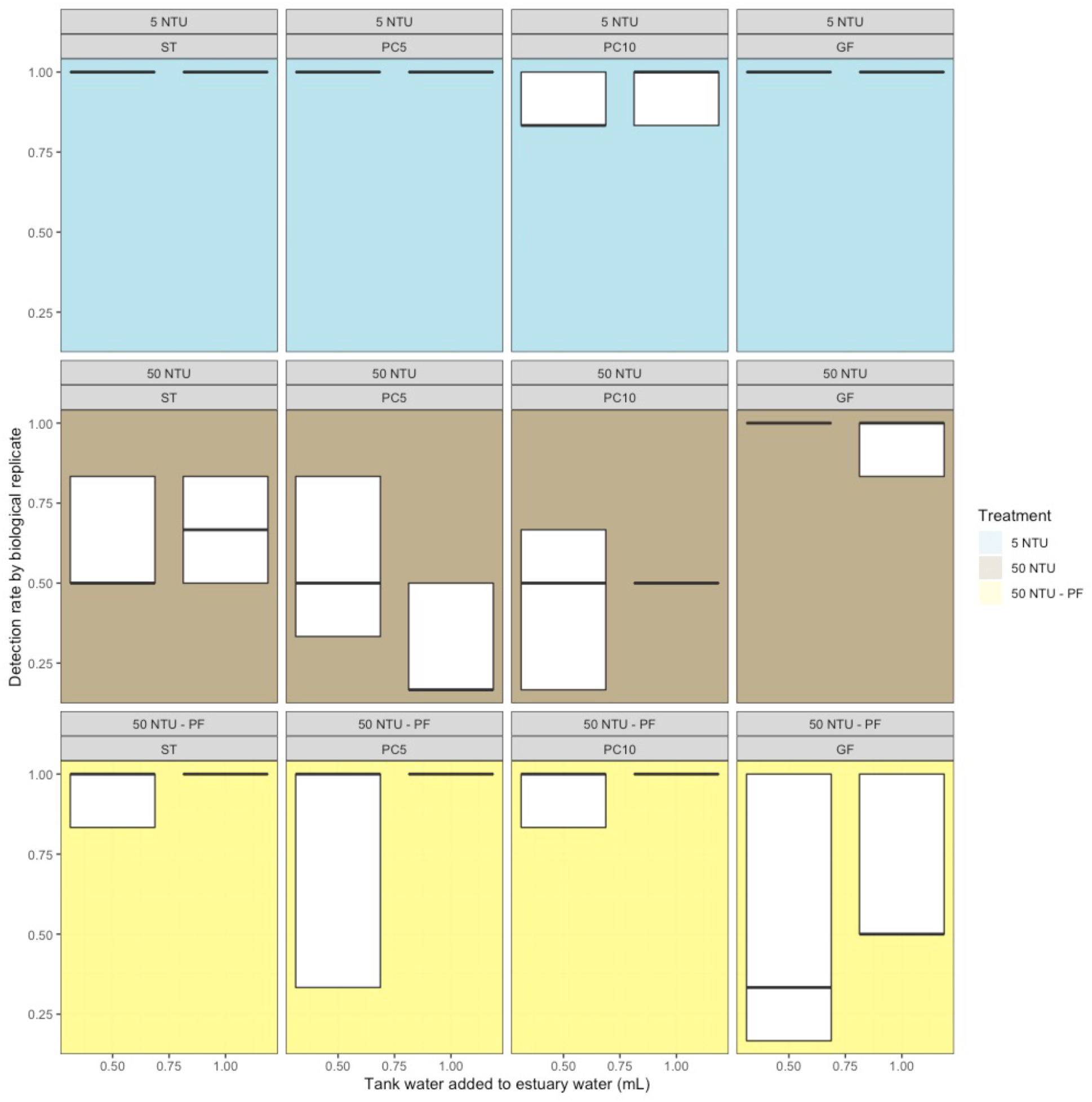
Results of filtration experiments as detection/non-detection of qPCR replicates (n=6) within 1-L biological replicates (n=3) for each treatment. Rows are treatment (turbidity and prefiltration), columns are filter type, and amount of delta smelt tank water added is within each box. GF, glass fiber filter; PC, polycarbonate filter; ST, Sterivex cartridge filter.

**Figure 5.**
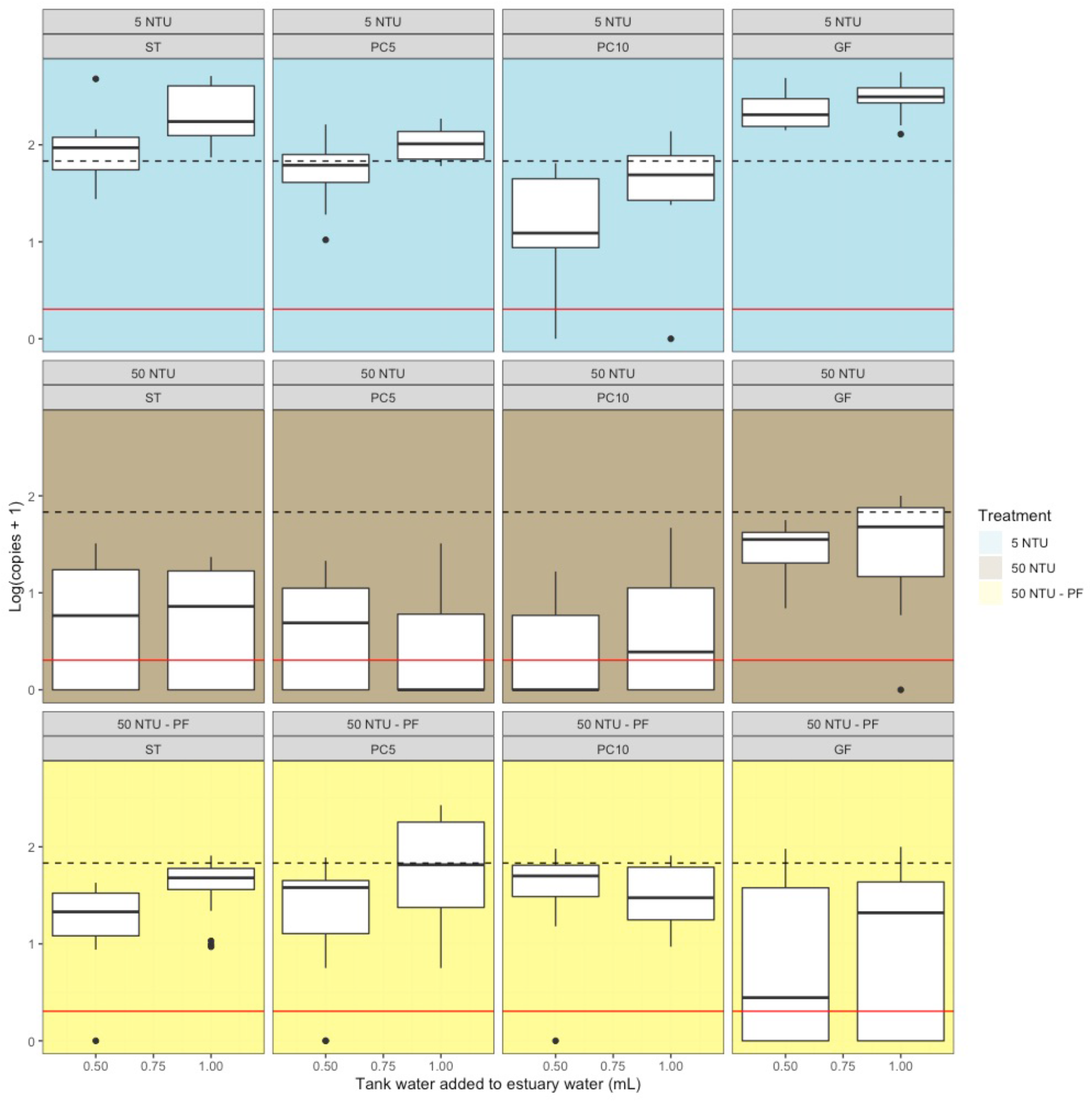
Results of filtration experiments as eDNA copies in qPCR replicates (n=6) within 1-L biological replicates (n=3) for each treatment. Rows are treatment (turbidity and prefiltration), columns are filter type, and amount of delta smelt tank water added is within each box. The black dotted line is the Limit of Quantification (LOQ) and the red dashed line is the Limit of Detection (LOD). GF, glass fiber filter; PC, polycarbonate filter; ST, Sterivex cartridge filter.

**Table 3.** Summary of detection rate and DNA copies detected for each treatment in the filtration experiment (turbidity, prefilter, and amount of tank water added (mL)). Further details in Supplementary Data S5. GF, glass fiber filter; PC, polycarbonate filter; ST, Sterivex PVDF filter.

| Treatment | Detection | DNA copies per reaction (6.1 µl template) |  |  |  |  | Approximate volume filtered (mL) | Mean DNA copies adjusted for volume filtered |
| --- | --- | --- | --- | --- | --- | --- | --- | --- |
|  | Positive qPCR replicates: 6 replicates x 3 bottles |  |  |  |  |  |  |  |
| Non-turbid |  | Mean | Median | SD | Min | Max |  |  |
| GF+500 | 18/18 | 233.4 | 203.7 | 94.3 | 141 | 489.8 | 1000 | 233.4 |
| PC10+500 | 16/18 | 24.2 | 11.4 | 22.8 | 0 | 63.8 | 1000 | 24.2 |
| PC5+500 | 18/18 | 68.6 | 60.3 | 46.1 | 9.6 | 161.4 | 1000 | 68.6 |
| ST+500 | 18/18 | 106.6 | 92.7 | 98.9 | 26.8 | 476.8 | 500 | 213.2 |
| GF+1000 | 18/18 | 324.7 | 310.1 | 111.9 | 128.4 | 560.4 | 1000 | 324.7 |
| PC10+1000 | 17/18 | 53.3 | 48.3 | 34.9 | 0 | 135.5 | 1000 | 53.3 |
| PC5+1000 | 18/18 | 106.4 | 103.5 | 40.4 | 58.8 | 187.1 | 1000 | 106.4 |
| ST+1000 | 18/18 | 238.1 | 173.3 | 157.2 | 73.4 | 506.6 | 500 | 476.2 |
| Turbid without prefilter |  |  |  |  |  |  |  |  |
| GF+500 | 18/18 | 30.8 | 34.7 | 15.8 | 5.9 | 55 | 1000 | 30.8 |
| PC10+500 | 8/18 | 3.3 | 0 | 4.7 | 0 | 15.7 | 450 | 7.3 |
| PC5+500 | 10/18 | 5.8 | 3.9 | 6.6 | 0 | 20.2 | 200 | 29 |
| ST+500 | 11/18 | 8.3 | 5 | 9.5 | 0 | 31 | 100 | 83 |
| GF+1000 | 17/18 | 46.9 | 47.4 | 33.7 | 0 | 99.4 | 1000 | 46.9 |
| PC10+1000 | 9/18 | 7.1 | 2.5 | 11.3 | 0 | 45.5 | 450 | 15.8 |
| PC5+1000 | 5/18 | 5.5 | 0 | 10 | 0 | 31.5 | 200 | 27.5 |
| ST+1000 | 12/18 | 8.4 | 6.5 | 8.3 | 0 | 22.5 | 100 | 84 |
| Turbid with prefilter |  |  |  |  |  |  |  |  |
| GF+500 | 9/18 | 23.2 | 3.3 | 34.9 | 0 | 94.2 | 1000 | 23.2 |
| PC10+500 | 17/18 | 49.3 | 49 | 27.2 | 0 | 96.5 | 750 | 65.7 |
| PC5+500 | 17/18 | 31.7 | 36.8 | 23.5 | 0 | 76.5 | 200 | 158.5 |
| ST+500 | 17/18 | 21.9 | 20.4 | 12.5 | 0 | 41.4 | 150 | 146 |
| GF+1000 | 9/18 | 29.3 | 19.9 | 33 | 0 | 98.7 | 1000 | 29.3 |
| PC10+1000 | 17/18 | 38.8 | 29.1 | 25.7 | 8.3 | 80.1 | 750 | 51.7 |
| PC5+1000 | 17/18 | 99.4 | 64.2 | 89.6 | 4.6 | 269.1 | 200 | 497 |
| ST+1000 | 17/18 | 44.7 | 46.8 | 22.2 | 8.4 | 80.7 | 150 | 298 |

### Statistical modeling

Model comparison did not support retaining a random effect term for individual bottles (Supplemental File S6). The highest weighted models tested for both response variables retained fixed effects filter type and prefilter, and an interaction between the filter type and prefilter (Table 4; Supplemental File S6). For eDNA copies as the response variable, the full model and a model where only filtration volume was missing. For detection/non-detection as a response variable, turbidity and tank water added were each missing from one of the top two models.

**Table 4.**
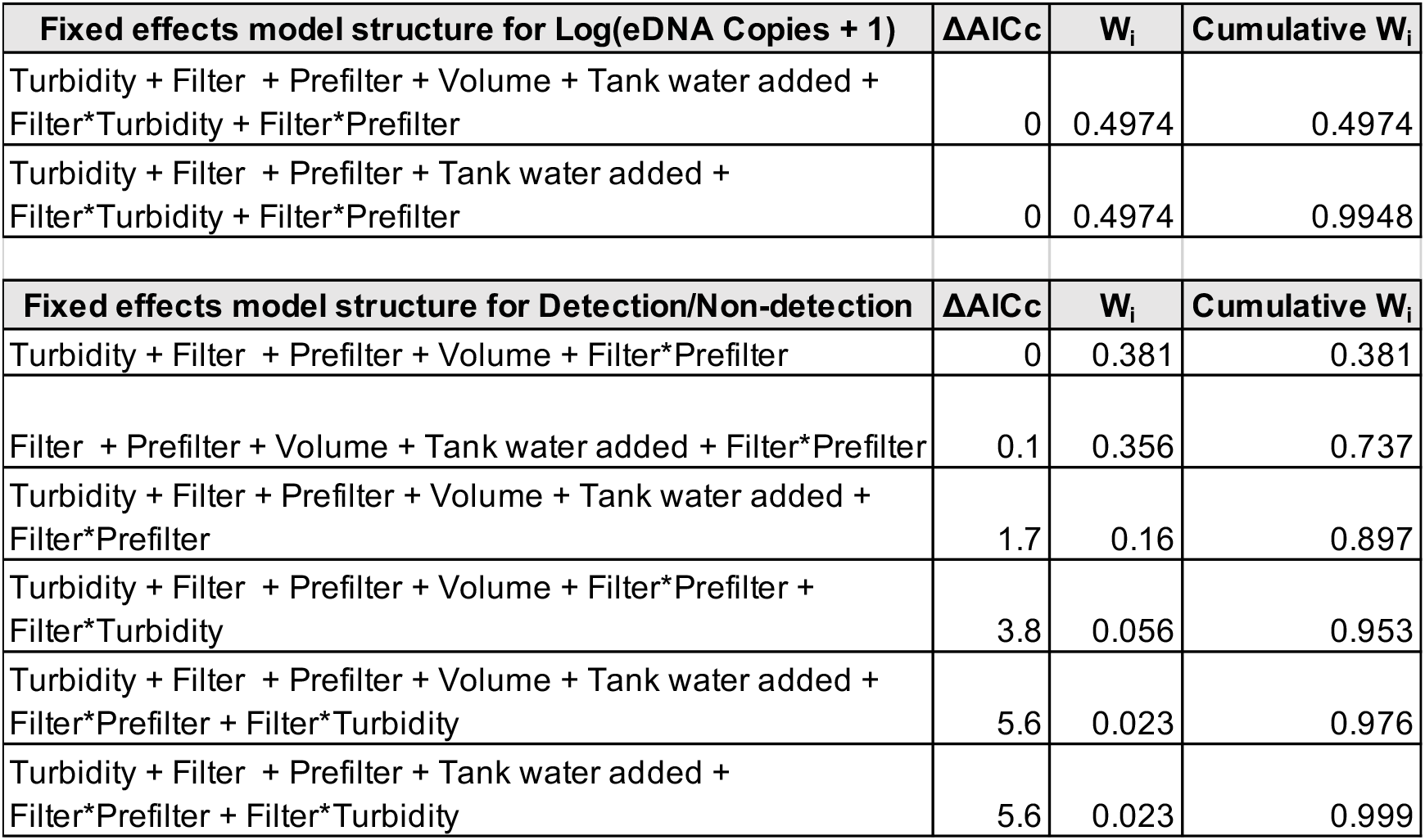
Summary of best models for success of delta smelt eDNA detection using (a) eDNA copies and (b) detection/non-detection as the response variables. Results of all models tested are provided Tables S6 and S7.

| **Fixed effects model structure for Log(eDNA Copies + 1)** | **$\Delta AIC_c$** | **$W_i$** | **Cumulative $W_i$** |
| --- | --- | --- | --- |
| Turbidity + Filter + Prefilter + Volume + Tank water added + Filter*Turbidity + Filter*Prefilter | 0 | 0.4974 | 0.4974 |
| Turbidity + Filter + Prefilter + Tank water added + Filter*Turbidity + Filter*Prefilter | 0 | 0.4974 | 0.9948 |
| **Fixed effects model structure for Detection/Non-detection** | **$\Delta AIC_c$** | **$W_i$** | **Cumulative $W_i$** |
| Turbidity + Filter + Prefilter + Volume + Filter*Prefilter | 0 | 0.381 | 0.381 |
| Filter + Prefilter + Volume + Tank water added + Filter*Prefilter | 0.1 | 0.356 | 0.737 |
| Turbidity + Filter + Prefilter + Volume + Tank water added + Filter*Prefilter | 1.7 | 0.16 | 0.897 |
| Turbidity + Filter + Prefilter + Volume + Filter*Prefilter + Filter*Turbidity | 3.8 | 0.056 | 0.953 |
| Turbidity + Filter + Prefilter + Volume + Tank water added + Filter*Prefilter + Filter*Turbidity | 5.6 | 0.023 | 0.976 |
| Turbidity + Filter + Prefilter + Tank water added + Filter*Prefilter + Filter*Turbidity | 5.6 | 0.023 | 0.999 |

## Discussion

In this study, we set out to untangle some of the challenges of eDNA detection of a very rare target organism in turbid conditions. First, we adapted a protocol for detection of delta smelt eDNA based on an assay developed for detection of delta smelt tissue in predator guts (Baerwald et al. 2011). The calculated Limit of Detection (LOD) and Limit of Quantification (LOD) indicate this part of the protocol is optimized for use eDNA detection of delta smelt. These limits can help guide interpretation of eDNA results (Figure 5).

We found that (1) turbidity decreased detection, (2) pore size appear less important than filter type for increasing detection in turbid water, and (3) prefiltration has mixed results. In non-turbid water, all filters except 10 μm polycarbonate filters had 100% detection and eDNA copies at or above the LOQ. This result is consistent with the presumed size of eDNA particles (1-10 μm; Turner et al., 2014). Filters with smaller pores (<1 μm) recommended to optimize eDNA capture can lead to reduced sample volumes and longer filtration times (Li et al. 2018). We did not see a clear pattern of larger pore sizes (as listed in filter description; Table 1) performing better in turbid water; filter material and construction may be more important characteristics. Despite low sample volumes, Sterivex cartridge filters used with a prefilter provided the most consistent results in terms of eDNA copy number. Cartridge filters are easier to protect from contamination in the field, however they are more expensive than the circular filters and extraction is time-consuming. Experiments indicated both a reduction in eDNA copies and detections in turbid water that could be partially mitigated with a prefilter for some filter types; prefilters did not appear to perform well when used with glass fiber filters. Filter type and prefilter status appeared to be the most important influences on detection. Glass fiber filters used without a prefilter provide appear to provide more efficient and economical detection of eDNA.

### Interpretation of low concentration eDNA

LOD and LOQ help establish standard practices for reporting eDNA detections, especially for detection of low-concentration eDNA (Klymus et al. 2019). The LOD of the Taqman assay used for delta smelt eDNA detection (Baerwald et al. 2011) was at the theoretical lowest limit of 3 copies per reaction, indicating that qPCR detection was well-optimized. eDNA copy numbers were generally below the LOQ (i.e., unreliable for quantification). In turbid water, there was not a clear distinction in detection between samples with 0.5mL and 1L of tank water added in turbid water (Figure 5). Copy number is likely an unreliable metric for using eDNA to model abundance or biomass of a rare fish in turbid water; presence/absence is a more straightforward signal to interpret when samples vary turbidity. These results also suggest that large turbidity differences between samples can prevent an apples-to-apples comparison of eDNA sample concentration for the same target species.

For rare species, even detection/non-detection can be challenging to interpret: when is a weak signal considered a positive detection? High Cq values (>40) are often treated as unreliable and therefore interpreted as potential false positive detections. However, the common practice of setting non-detect values to 40 may introduce bias (McCall et al. 2014) and increase the false negative detection rate. As in many areas of science, it is impossible to “prove the negative.” In this study, 3 of 432 qPCR reactions (less than 1%) assaying samples with the addition of delta smelt tank water generated Cq values >40 (Table S4). The results of our analysis of these samples known to contain target eDNA suggest that, although rare, Cq values >40 can represent true detections. There was no evidence of contamination in negative control samples. In addition, although the protocol ran 50 cycles, there were no detections above Cq of 42. Similarly, an evaluation of eDNA metabarcoding laboratory protocols shows that, above a certain threshold, additional PCR cycles do not improve species detection (Stoeckle et al. 2022). Given the current limits of technology, interpretation of weak signals is a balancing act between signal and noise and likely specific to each particular application of eDNA.

### Effect of suspended particulate matter on filtering and detection

As expected, filters with smaller pores filtered less water. A positive relationship has been demonstrated between sample volume and species detection in both turbid (Williams et al. 2017) and non-turbid water, (Wilcox et al. 2015; Wilcox et al. 2016; Sepulvida et al. 2019; Bedwell and Goldberg 2020), although under certain conditions there may not be a relationship between sample volume and species detection (Mächler et al. 2016). Suspended particulate matter clogs filters and increases filtration pressure, requiring re-optimization of the capture method (Thomas et al. 2018). Although we did not measure membrane pressure during our experiment, our data is consistent with poor eDNA capture due to high membrane pressure. Membrane pressure may decrease eDNA retention by breaking apart clumps of cells or bursting cells or mitochondria (Thomas et al. 2018). Glass fiber filters, which performed best in turbid water, have a completely different pore type and construction than the other filters used in this study (Table 1).

Pore sizes are not necessarily comparable between filter materials. Filter materials have different pore types. Absolute pores (e.g., polycarbonate and polyvinylidene fluoride (PVDF)) are uniform in size. These filters act as a screen, retaining all particles larger than the pores on the filter surface. Glass fiber filters have nominal pores that are irregular and retain only a percentage of particles larger than the pore size and are depth filters with multiple layers that trap particles inside a structure. (Cellulose and cellulose nitrate are other nominal pore filter types that are commonly used for eDNA capture.) Despite lower capture efficiency, the thickness of depth filters may provide relatively more space to capture particles in turbid water. Glass fiber filters are significantly less expensive than the other filters used in this study. However, PowerWater extractions use a bead beating step that causes glass fiber to become sponge-like, requiring more time and care to separate the supernatant from the beads and filter.

Finally, eDNA interactions with turbidity may vary depending characteristics of the particulates. In samples from experimental ponds with turbidity up to 60 NTU, turbidity was positively associated with eDNA detected using 10-μm pore polycarbonate filters and prefiltration, suggesting that eDNA was sticking to larger phytoplankton in the ponds (Barnes et al. 2020). Turbidity in river-dominated estuaries like the SFE is mainly caused by river inputs of suspended particulate matter and resuspension of bottom sediments rather than phytoplankton (Cloern 1987). While eDNA can be detected in samples of suspended particulate matter collected in sedimentation boxes in rivers (Díaz et al. 2020), it is not clear if eDNA co-occurs with these particulates or is stuck to them. In our experiment, turbidity reduced detection rates and we did not see evidence of eDNA sticking to particulates. Cellular studies suggest that cells may be more likely to adhere to each other than foreign material (Coman 1961).

### Effect of prefiltration on detection

Prefiltration is sometimes recommended to increase sample volume and decrease filtration time. Our results indicated a significant interaction between filter type and prefiltration. One explanation for the negative impact of prefiltration on glass fiber filters is that prefiltration breaks eDNA particles that cannot be efficiently captured by glass fiber. For example, clumps of cells may be broken up into individual cells or whole cells may be reduced to mitochondria. In cases where copy number quantification is feasible, prefiltration in combination with end filters with small pores may provide more consistent results across turbidity conditions. Prefiltration also increases the cost and effort in filtering. If the study goal is to determine species presence/absence, and budget or time is limited, then our results indicate the most practical approach for rare species detection in turbid conditions is to use glass fiber filters without prefiltration.

### Future directions

Experimental work is necessarily limited to a relatively narrow range of treatments but can help tease out the effects of turbidity on eDNA detection without the complexity of natural systems. eDNA is not homogeneously distributed in natural environments, making it more difficult to determine if decreased detections in turbid water reflect real patterns of occurrence or limitations of eDNA detection. eDNA samples collected in relatively more turbid water (Secchi depth 73 cm; see Supplemental File S7 for the relationship between Secchi depth and turbidity) can yield more detections when turbid conditions are more favorable (Kumar et al. 2021). Experimental work encompassing a wide range of turbidity conditions could help further our understanding of eDNA detection in turbid water. A correction factor could be developed to account for the decrease in detection observed as turbidity increases, although such corrections may be particle- or habitat-specific.

Finally, turbidity (an optical measurement) is often approximated using other metrics (e.g., water clarity, total suspended solids, filtration time), severely restricting the ability to compare eDNA studies conducted in turbid water. Comparable measurements may help optimize eDNA methods and data interpretation when turbidity is a significant characteristic of the target species habitat. Optical measurements taken using turbidimeters and probes are the most objective, repeatable, and accurate across a broad range of values of turbidity and can be used in most water bodies (Pickering 1976). Water clarity measured by Secchi disk is a less expensive alternative but cannot be used certain conditions (e.g., fast moving water) and is subject to human error (Carlson and Simpson 1996). Filtration time as a proxy for turbidity is less useful because it is difficult to calibrate across different studies and filtration set-ups.

## Conclusions

More knowledge of endangered fishes is needed to meet the conservation goals (Guy et al. 2021), presenting a perfect opportunity to employ high sensitivity methods like eDNA detection for surveys and monitoring. eDNA detection methods, however, are not “one size fits all” (Barnes and Turner 2016; Kumar et al. 2021). We use this comparison of eDNA capture methods under controlled conditions to help guide best practices for the real-world challenge of detecting a rare species in turbid conditions. Turbidity and filter type influence eDNA detection success and prefiltration may not always be beneficial. These findings provide optimism that reliable and repeatable eDNA detections of rare species are possible in turbid when appropriately optimized methods are used.

## Data accessibility

Data and R code are available at https://github.com/annholmes/eDNA-experiments-in-turbid-water.

## Acknowledgements

Thank you to Grace Auringer, Alyssa Benjamin, Alisha Goodbla, Leslie Guerrero, and Shannon Kieran for assistance and advice on sampling and laboratory work, to Dennis Cocherell, Brittany Davis, Luke Ellison, Nann Fangue, and Tien-Chieh Hung for access to delta smelt tanks, and to Ted Sommer and Andrea Schreier for feedback and support. We are grateful to Chris Hart, Jessica Adams, Denise Barnard, Bill Powell, and the US Fish and Wildlife Service Enhanced Delta Smelt Monitoring (EDSM) Program for facilitating paired eDNA sampling with trawl surveys. The University of California, Davis sits on Patwin Land.

## Author contributions

Conception and design: AH, MB, JR, BS, AF

Experiments and laboratory work: AH

Field sampling: AH, BM

Data analysis: AH with input from BM

Prepared first draft of manuscript: AH

Revised and approved: AH, MB, JR, BS, BM, AF

