## Supplementary File 1 for "Experimental evaluation of environmental DNA detection of a rare fish in turbid water"

**Supplemental File 2: Detailed extraction protocol for glass fiber filters**

Qiagen Experienced User Protocol in regular text
***Modifications and notes from this study in bold italics***

Important points before starting

- Solution PW1 must be warmed to 55°C ***(or 65°C)*** for 5–10 min to dissolve precipitates prior to use. Solution PW1 should be used while still warm.
- If Solution PW3 has precipitate, heat to 55°C for 5–10 min to dissolve precipitate.
- Shake to mix Solution PW4 before use.
- Perform all centrifugation steps at room temperature (15–25°C).
- ***Preheat incubator to 65ºC.***

**Procedure**
***Steps 1-3 pertain to filtration and filter preservation; see main text for the methods used in this study. This protocol works best for extracting DNA from 15-20 filters at a time.***

Step 4. ***Remove tubes from the freezer and allow the filters to thaw. Using sterile forceps, remove each filter from the storage tube and*** insert each filter into a 5 ml PowerWater Bead Pro Tube. ***Using 1-2 sterile forceps, unfold the filter inside the 5 ml tube and expose the side with captured material. This can be a tricky procedure and proper folding of filters after fiiltering greatly improves this process (see main text).***

Step 5. Add 1 ml of ***warm*** Solution PW1 to ***filter surface inside*** the PowerWater Bead Pro Tube.

Note: For samples containing organisms that are difficult to lyse (e.g., fungi and algae) an additional heating step can be included. See Alternative Lysis Methods in the Troubleshooting Guide. ***Incubate for 10 min at 65ºC.***

Step 6. ***Tighten caps on 5 ml tubes after heat incubation.*** Secure the tube horizontally to a Vortex Adapter (cat. no. 13000-V1-5 or 13000-V1-15).

Step 7. Vortex at maximum speed for 5 min. ***(Centrifuge was not used in this protocol because the equipment was not available.)***

***Notes: During this step the glass fiber filter should fully break apart. Sometimes the supernatant gets foamy. Remove tubes from vortexer and stand upright for foam to settle.***

Step 8. Transfer the supernatant to a clean 2 ml collection tube (provided). Draw up the supernatant using a 1 ml pipette tip by placing it down into the beads. Note: Placing the pipette tip down into the beads is required. Pipette until you have removed all the supernatant.

***Notes: Qiagen protocol says expect to recover 600–650 µl of supernatant. As glass fiber filters hold liquid, the amount of supernatant recovered is generally 600-1200 µl. For turbid samples, the supernatant should reflect the color of the filter (usually greenish or brownish).***

Step 9. Centrifuge at 13,000 x g (~8000 rpm) for 1 min.

Step 10. Avoiding the pellet, transfer the supernatant to a clean 2 ml collection tube (provided).

Step 11. Add 200 µl of Solution IRS and vortex briefly to mix. Incubate at 2–8°C for 5 min. ***(Longer incubation times appear to have no effect on extraction efficiency; this incubation step is a good time to break, if needed.)***

Step 12. Centrifuge the tubes at 13,000 x g (~8000 rpm) for 1 min.

Step 13. Avoiding the pellet, transfer the supernatant to a clean 2 ml collection tube (provided).

Step 14. Add 650 µl of Solution PW3 and vortex briefly to mix. ***If the volume of the supernatant and PW3 exceeds the capacity of the 2 ml tube, split the supernatant into two tubes and add PW3 to the second tube.***

Step 15. Load 650 µl of supernatant onto an MB Spin Column. Centrifuge at 13,000 x g (~8000 rpm) for 1 min. Discard the flow-through. Repeat until all the supernatant has been processed.

Step 16. Place the MB Spin Column Filter into a clean 2 ml collection tube (provided). ***Use the provided 1.5 ml collection tubes but manually remove the caps to prevent them from coming off in the centrifuge.***

Step 17. Add 650 µl of Solution PW4 (shake before use). Centrifuge at 13,000 x g (~8000 rpm) for 1 min.

Step 18. ***Place sprin column into a clean 2 ml collection tube,*** add 650 µl of ethanol (provided) and centrifuge at 13,000 x g (~8000 rpm) for 1 min. ***The bottom of the spin column will likely be in contact with the flow-through if using the collection tubes provided; discard the flow-through and centrifuge again at 13,000 x g (~8000 rpm) for 1 min.***

Step 19. Discard the flow-through and centrifuge again at 13,000 x g (~8000 rpm) for 2 min.

Step 20. Place the MB Spin Column into a clean 2 ml collection tube (provided).

***Note: Use Eppendorf 1.5 ml DNA LoBind tubes instead of collection tubes provided in kit.***

Step 21. Add 100 µl of Solution EB to the center of the white filter membrane. ***Solution should be added carefully so that it sits on the membrane, not the side of the tube. Incubate for at least 20 min to increase DNA yield.***

Step 22. Centrifuge at 13,000 x g (~8000 rpm) for 1 min.

Step 23. Discard the MB Spin Column. The DNA is now ready for downstream applications. Note: We recommend storing DNA frozen (–90°C to –15°C) as Solution EB does not contain EDTA. To concentrate DNA, see the Troubleshooting Guide.

***Note: eDNA should be stored in LoBind tubes.***
