## Supplementary File 2 for "Experimental evaluation of environmental DNA detection of a rare fish in turbid water"

### Supplementary File 1

Delta smelt eDNA field sampling paired with Enhanced Delta Smelt Monitoring (EDSM) kodiak trawl survey. Trawl data (DSM trawl) reports the number of individual delta smelt captured in each trawl. Delta smelt eDNA detections (DSM eDNA) are shown as the proportion of positive qPCR reactions out of 8 total reactions.

| Date | Gear In | Station | Tow # | Lat | Long | DSM trawl | DSM eDNA | Turb (NTU) | ºC | Sp cnd | DO | Current | Tide prev | Tide next | Tide dir | Weather |
|---|---|---|---|---|---|---|---|---|---|---|---|---|---|---|---|---|
| 1/3/17 | 0939 | HB4 | 1 | 38.0750 | -121.9310 | 0 | 0/8 | 80 | 8.0 | 166.0 | 10.6 | -1.37 | 924 | 1448 | ebb | rain |
| 1/3/17 | 0959 | HB4 | 2 | 38.0730 | -121.9580 | 0 | 0/8 | 70.8 | 8.1 | 171.0 | 10.4 | -1.37 | 924 | 1448 | ebb | rain |
| 1/3/17 | 0808 | HB5 | 1 | 38.0477 | -121.9304 | 0 | 0/8 | 46.8 | 8.1 | 188.0 | 10.7 | 1.72 | 254 | 924 | flood | rain |
| 1/3/17 | 0828 | HB5 | 2 | 38.0477 | -121.9304 | 0 | 0/8 | 51.2 | 8.1 | 140.0 | 10.5 | 1.72 | 254 | 924 | flood | rain |
| 1/3/17 | 1136 | RV3 | 1 | 38.0897 | -121.7377 | 0 | 0/8 | 41.6 | 8.3 | 85.5 | 10.6 | -1.37 | 924 | 1448 | ebb | rain |
| 1/3/17 | 1203 | RV3 | 2 | 38.0890 | -121.7372 | 0 | 0/8 | 29.3 | 8.2 | 105.4 | 10.3 | -1.37 | 924 | 1448 | ebb | rain |
| 1/3/17 | 1217 | RV3 | 3 | 38.0894 | -121.7366 | 0 | 0/8 | 32.4 | 8.3 | 61.9 | 10.5 | -1.37 | 924 | 1448 | ebb | rain |
| 1/3/17 | 1227 | RV3 | 4 | 38.0897 | -121.7372 | 0 | 0/8 | 30.5 | 8.3 | 102.7 | 10.4 | -1.37 | 924 | 1448 | ebb | rain |
| 1/6/17 | 0937 | HB6 | 1 | 38.0415 | -121.9081 | 0 | 0/8 | 59.9 | 8.2 | 127.2 | 9.6 | 1.78 | 506 | 1130 | flood | clear |
| 1/6/17 | 1014 | HB6 | 4 | 38.0427 | -121.9118 | 1 | 1/8 | 67.8 | 8.2 | 160.0 | 9.3 | 1.78 | 506 | 1130 | flood | clear |
| 1/6/17 | 0823 | LSR13 | 1 | 38.0217 | -121.8110 | 2 | 1/8 | 50.4 | 8.2 | 59.3 | 9.3 | 1.78 | 506 | 1130 | flood | cloudy |
| 1/6/17 | 0842 | LSR13 | 2 | 38.0222 | -121.8113 | 3 | 1/8 | 53.5 | 8.2 | 148.7 | 9.6 | 1.78 | 506 | 1130 | flood | clear |
| 1/24/17 | 0932 | HB52 | 1 | 38.0479 | -121.9305 | 0 | 0/8 | 149 | 9.1 | 96.2 | 10.9 | 1.48 | 754 | 1412 | flood | clear |
| 1/24/17 | 0944 | HB52 | 2 | 38.0473 | -121.9272 | 0 | 0/8 | 144 | 9.1 | 97.0 | 10.8 | 1.48 | 754 | 1412 | flood | clear |
| 1/24/17 | 1025 | HB52 | 5 | 38.0470 | -121.9248 | 8 | 1/8 | 145 | 9.1 | 100.3 | 10.6 | 1.48 | 754 | 1412 | flood | clear |
| 1/24/17 | 1136 | RV51 | 1 | 38.1289 | -121.6918 | 0 | 0/8 | 112 | 8.9 | 77.5 | 10.9 | 1.48 | 754 | 1412 | flood | cloudy |
| 1/24/17 | 1147 | RV51 | 2 | 38.1294 | -121.6913 | 0 | 0/8 | 124 | 8.9 | 77.9 | 11.1 | 1.48 | 754 | 1412 | flood | cloudy |

| | | | | | | | | | | | | | | | | |
|---|---|---|---|---|---|---|---|---|---|---|---|---|---|---|---|---|
| 1/24/17 | 1158 | RV51 | 3 | 38.1301 | -121.6908 | 0 | 0/8 | 119 | 8.9 | 76.2 | 11.2 | 1.48 | 754 | 1412 | flood | cloudy |
| 1/24/17 | 1208 | RV51 | 4 | 38.1306 | -121.6904 | 0 | 0/8 | 118 | 8.9 | 76.7 | 11.2 | 1.48 | 754 | 1412 | flood | cloudy |
| 1/24/17 | 0822 | SBM53 | 1 | 38.0835 | -121.9945 | 0 | 0/8 | 118 | 9.0 | 122.9 | 10.7 | 1.48 | 754 | 1412 | flood | cloudy |
| 1/24/17 | 0833 | SBM53 | 2 | 38.0829 | -121.9929 | 0 | 0/8 | 123 | 9.0 | 126.2 | 11 | 1.48 | 754 | 1412 | flood | cloudy |
| 1/24/17 | 0844 | SBM53 | 3 | 38.0841 | -121.9898 | 0 | 0/8 | 151 | 9.0 | 127.7 | 10.9 | 1.48 | 754 | 1412 | flood | cloudy |
| 1/24/17 | 0857 | SBM53 | 4 | 38.0843 | -121.9871 | 1 | 1/8 | 142 | 9.0 | 121.9 | 10.6 | 1.48 | 754 | 1412 | flood | clear |
| 2/9/17 | 1220 | RV51 | 5 | 38.1311 | -121.6897 | 0 | 0/8 | 126 | 9.0 | 77.2 | 11.2 | 1.48 | 754 | 1412 | flood | cloudy |
| 2/9/17 | 0821 | SBM60 | 1 | 38.0796 | -121.9819 | 0 | 0/8 | 84.6 | 11.5 | 99.4 | 13.2 | 4.48 | 729 | 1305 | flood | cloudy |
| 2/9/17 | 0831 | SBM60 | 2 | 38.0794 | -121.9823 | 0 | 0/8 | 100 | 11.5 | 101.8 | 12.9 | 4.48 | 729 | 1305 | flood | cloudy |
| 2/9/17 | 0852 | SBM60 | 3 | 38.0790 | -121.9815 | 0 | 0/8 | 103 | 11.6 | 110.1 | 12.8 | 4.48 | 729 | 1305 | flood | cloudy |
| 2/9/17 | 0904 | SBM60 | 4 | 38.0783 | -121.9807 | 0 | 0/8 | 115 | 11.9 | 111.5 | 12.6 | 4.48 | 729 | 1305 | flood | cloudy |
| 2/9/17 | 0916 | SBM60 | 5 | 38.0777 | -121.9804 | 0 | 0/8 | 114 | 11.9 | 113.3 | 12.5 | 4.48 | 729 | 1305 | flood | cloudy |
| 2/9/17 | 0958 | SBM62 | 1 | 38.0957 | -122.0312 | 0 | 0/8 | 94.8 | 11.4 | 110.3 | 12.9 | 4.48 | 729 | 1305 | flood | cloudy |
| 2/9/17 | 1009 | SBM62 | 2 | 38.0935 | -122.0257 | 0 | 0/8 | 77.7 | 11.6 | 114.8 | 12.9 | 4.48 | 729 | 1305 | flood | rain |
| 2/9/17 | 1031 | SBM62 | 3 | 38.0951 | -122.0331 | 1 | 1/8 | 79.2 | 11.5 | 106.0 | 12.8 | 4.48 | 729 | 1305 | flood | rain |
