## Supplementary File 4 for "Experimental evaluation of environmental DNA detection of a rare fish in turbid water"

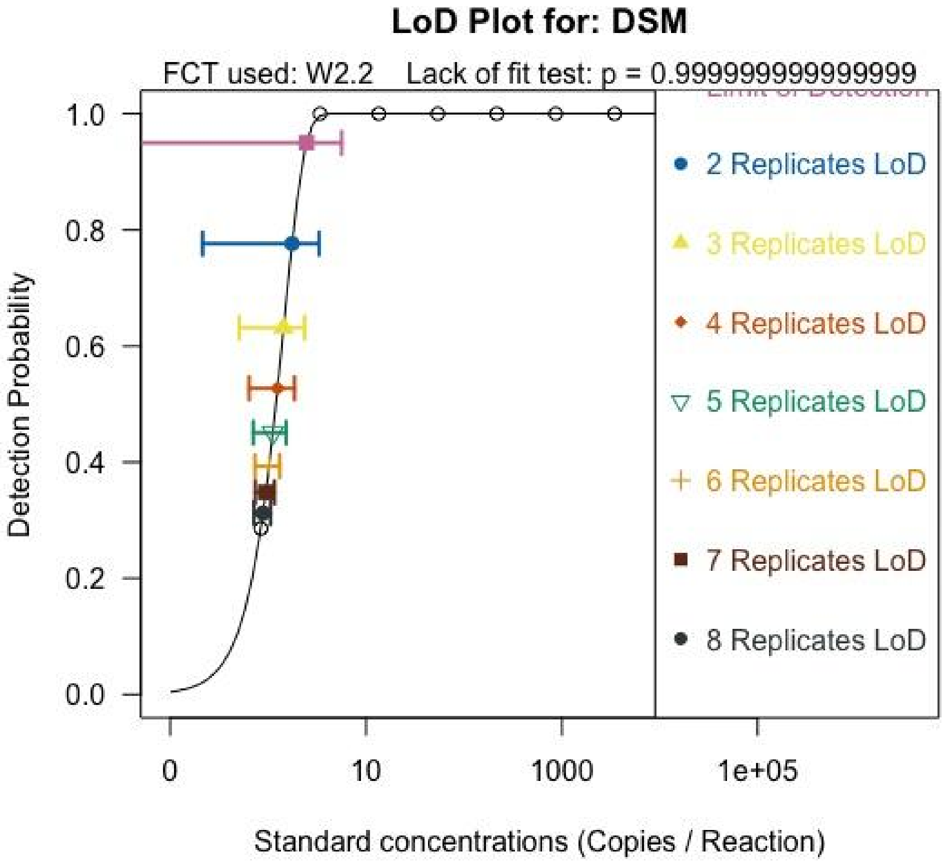


| Estimate | Std.Error | Lower | Upper | LOD | Assay |
|---|---|---|---|---|---|
| 2.465 | 1.589 | -0.690 | 5.620 | 1rep.LOD | DSM |
| 1.752 | 0.775 | 0.213 | 3.291 | 2rep.LOD | DSM |
| 1.435 | 0.468 | 0.506 | 2.363 | 3rep.LOD | DSM |
| 1.245 | 0.306 | 0.638 | 1.852 | 4rep.LOD | DSM |
| 1.116 | 0.208 | 0.703 | 1.528 | 5rep.LOD | DSM |
| 1.020 | 0.145 | 0.731 | 1.308 | 6rep.LOD | DSM |
| 0.945 | 0.107 | 0.734 | 1.157 | 7rep.LOD | DSM |
| 0.885 | 0.086 | 0.714 | 1.056 | 8rep.LOD | DSM |

Effective Limit of Detection (LOD) plot and related data showing detection probability and 95% confidence intervals for standard concentrations assayed in 1 to 8 qPCR replicates calculated using standardized methods for eDNA analysis (Klymus et al. 2019; Merkes et al. 2019).

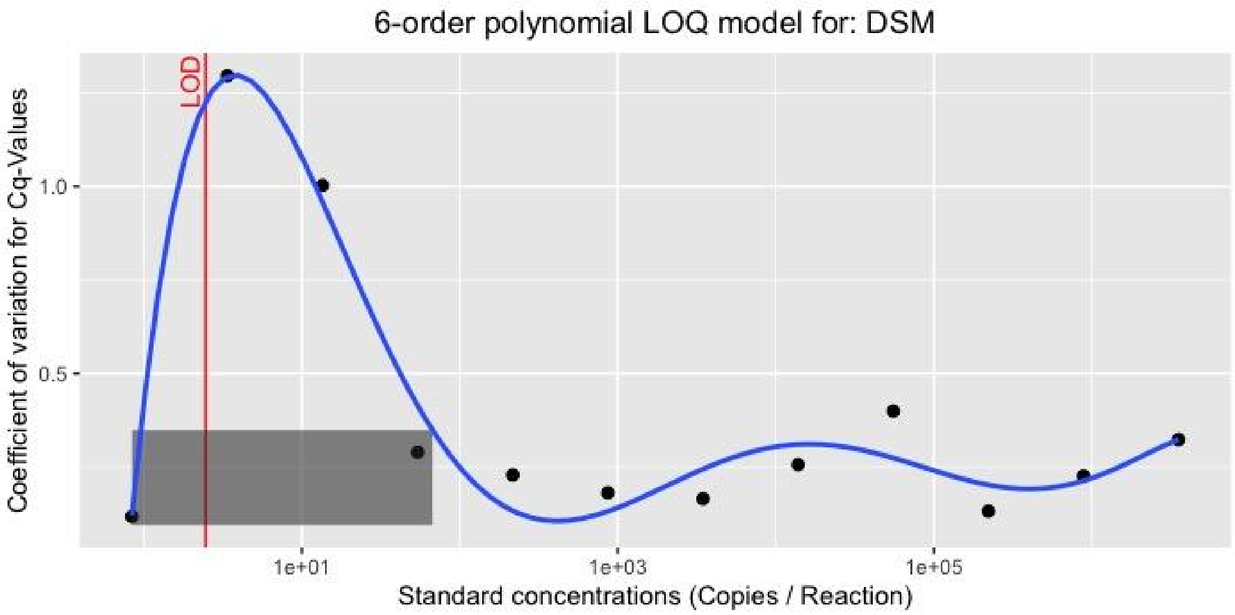


The 6-order polynomial Limit of Quantification (LOQ) model provides a visualization of the concentration where assay precision begins to decrease, i.e. the intersection of the blue LOQ line and the top right corner of the gray box. LOQ is defined as the lowest concentration at which the CV of qPCR results is less than 35% and the LOD (red line) is provided for reference. The model was calculated, visualized and interpreted using standardized methods for eDNA analysis (Klymus et al. 2019; Merkes et al. 2019).
