## Supplementary File 6 for "Experimental evaluation of environmental DNA detection of a rare fish in turbid water"

**Supplementary File 5**

Comparison of models with and without a random effect for individual bottles (i.e biological replicates) for delta smelt eDNA detection using (a) eDNA copies and (b) detection/non-detection as the response variables.

a.

| Fixed effects model structure for Log(eDNA Copies + 1) | Random effect | ΔAICc | $W_i$ |
|---|---|---|---|
| Turbidity + Filter + Prefilter + Volume + Tank water added + Filter*Turbidity + Filter*Prefilter | - | 0 | 1 |
| Turbidity + Filter + Prefilter + Volume + Tank water added + Filter*Turbidity + Filter*Prefilter | Bottle | 68.3 | <0.001 |

b.

| Fixed effects model structure for Detection/Non-detection | Random effect | ΔAICc | $W_i$ |
|---|---|---|---|
| Turbidity + Filter + Prefilter + Volume + Tank water added + Filter*Turbidity + Filter*Prefilter | - | 0 | 1 |
| Turbidity + Filter + Prefilter + Volume + Tank water added + Filter*Turbidity + Filter*Prefilter | Bottle | 91 | <0.001 |

Results of all models tested. A summary of best models is provided in Table 4.

a.

| Fixed effects model structure for Log(eDNA Copies + 1) | ΔAICc | $W_i$ |
|---|---|---|
| Turbidity + Filter + Prefilter + Volume + Tank water added + Filter*Turbidity + Filter*Prefilter | 0 | 0.4974 |
| Turbidity + Filter + Prefilter + Tank water added + Filter*Turbidity + Filter*Prefilter | 0 | 0.4974 |
| Turbidity + Filter + Prefilter + Volume + Tank water added + Filter_type*Prefilter | 9.9 | <0.001 |
| Turbidity + Filter + Prefilter + Tank_water + Filter*Prefilter | 12.2 | <0.001 |
| Turbidity + Filter + Prefilter + Volume + Filter*Turbidity + Filter*Prefilter | 14.1 | <0.001 |
| Turbidity + Filter + Prefilter + Volume + Filter*Prefilter | 23.5 | <0.001 |
| Filter + Prefilter + Volume + Tank water added + Filter*Prefilter | 89.7 | <0.001 |
| Turbidity + Filter + Prefilter + Volume + Tank water added + Filter*Turbidity | 95.8 | <0.001 |
| Turbidity + Filter + Prefilter + Volume + Filter*Turbidity | 106.6 | <0.001 |
| Turbidity + Filter + Volume + Tank water added + Filter*Turbidity | 114.5 | <0.001 |
| Turbidity + Filter + Prefilter + Tank water added + Filter*Turbidity | 116.8 | <0.001 |
| Turbidity + Filter + Prefilter + Tank water added | 142.3 | <0.001 |
| Turbidity + Filter + Prefilter + Volume + Tank water added | 144.2 | <0.001 |
| Turbidity + Filter + Prefilter + Volume | 153.5 | <0.001 |
| Turbidity + Prefilter + Volume + Tank water added | 171.9 | <0.001 |
| Turbidity + Filter + Volume + Tank water added | 207.3 | <0.001 |
| Filter + Prefilter + Volume + Tank water added | 252.3 | <0.001 |
| Intercept only | 385.6 | <0.001 |
| | Total | 1 |

b.

| Fixed effects model structure for Detection/Non-detection | ΔAICc | $W_i$ |
|---|---|---|
| Turbidity + Filter + Prefilter + Volume + Filter*Prefilter | 0 | 0.381 |
| Filter + Prefilter + Volume + Tank water added + Filter*Prefilter | 0.1 | 0.356 |
| Turbidity + Filter + Prefilter + Volume + Tank water added + Filter*Prefilter | 1.7 | 0.160 |
| Turbidity + Filter + Prefilter + Volume + Filter*Prefilter + Filter*Turbidity | 3.8 | 0.056 |
| Turbidity + Filter + Prefilter + Volume + Tank water added + Filter*Prefilter + Filter*Turbidity | 5.6 | 0.023 |
| Turbidity + Filter + Prefilter + Tank water added + Filter*Prefilter + Filter*Turbidity | 5.6 | 0.023 |
| Turbidity + Filter + Prefilter + Volume + Tank water added + Filter_type*Prefilter | 28.9 | <0.001 |
| Turbidity + Filter + Prefilter + Volume + Filter*Turbidity | 74.8 | <0.001 |
| Turbidity + Filter + Prefilter + Volume + Tank water added + Filter*Turbidity | 76.6 | <0.001 |
| Turbidity + Filter + Volume + Tank water added + Filter*Turbidity | 81.1 | <0.001 |
| Turbidity + Filter + Prefilter + Volume | 81.3 | <0.001 |
| Turbidity + Filter + Prefilter + Volume + Tank water added | 83.1 | <0.001 |
| Turbidity + Filter + Prefilter + Tank water added | 86.5 | <0.001 |
| Turbidity + Prefilter + Volume + Tank water added | 86.7 | <0.001 |
| Turbidity + Filter + Prefilter + Tank water added + Filter*Turbidity | 90.7 | <0.001 |
| Filter + Prefilter + Volume + Tank water added | 93.6 | <0.001 |
| Turbidity + Filter + Volume + Tank water added | 107.3 | <0.001 |
| Intercept only | 150.2 | <0.001 |
| | Total | 1 |
