## Supplementary File 7 for "Experimental evaluation of environmental DNA detection of a rare fish in turbid water"

**Supplementary File 6**

Turbidity and water clarity

Although optical turbidity (nephelometry or optical scattering) and Secchi depth (water clarity or transparency) are not equivalent metrics, both are used to measure “turbidity.” The main advantages of Secchi depth over turbidimeters are low cost and simplicity of the instrument. The disadvantages of Secchi depth are the influences of human error and sunlight (Carlson and Simpson 1996). Secchi depth also must be taken in situ. Secchi depth is used widely and it is generally viewed as a “good enough” means to approximate turbidity (Pickering 1976). However, there is no easy way to convert Secchi depth to optical turbidity units because their precise relationship varies by habitat. In the case of a Secchi depth measurement of 73 cm (Kumar et al. 2021), conversion to optical units using data from another system (USGS, 2016) yields a turbidity measurement of ~10 NTU. Thus, the “high turbidity” water in this study is more similar to our study’s non-turbid water (5 NTU) than turbid water (50 NTU). As discussed in the paper, “high turbidity” is a relative measurement and the definition of “high” justifiably varies greatly across different habitats and ecosystems. The concurrent use of two metrics with an imprecise relationship in the literature makes it more difficult to assess the impact of turbidity on eDNA detection in real-world scenarios.
